# Application of Parallel Reaction Monitoring in ^15^N labeled Samples for Quantification

**DOI:** 10.1101/2021.12.01.470846

**Authors:** Andres V. Reyes, Ruben Shrestha, Peter R. Baker, Robert J. Chalkley, Shou-Ling Xu

## Abstract

Accurate relative quantification is critical in proteomic studies. The incorporation of stable isotope ^15^N to plant-expressed proteins *in vivo* is a powerful tool for accurate quantification with a major advantage of reducing preparative and analytical variabilities. However, ^15^N labeling quantification has several challenges. Less identifications are often observed in the heavy labeled samples because of incomplete labeling, resulting in missing values in reciprocal labeling experiments. Inaccurate quantification can happen when there is contamination from co-eluting peptides or chemical noise in the MS^1^ survey scan. These drawbacks in quantification can be more pronounced in less abundant but biologically interesting proteins, which often have very few identified peptides. Here we demonstrate the application of parallel reaction monitoring (PRM) to ^15^N labeled samples on a high resolution, high mass accuracy Orbitrap mass spectrometer to achieve reliable quantification even of low abundance proteins in samples.

## 1.1 Introduction

Relative quantitative comparisons between biological samples are critical in proteomic studies to uncover key differences of interest. The incorporation of stable isotope ^15^N to labeled proteins in plants is one powerful strategy that can be used to achieve such information. In such a quantitative experiment, one sample contains the most common light isotope (^14^N), and the other is labeled with a stable heavy isotope (^15^N) in the form of nitrogen salts, such as ^15^NH_4_^15^NO_3_, K^15^NO_3_, or ^15^NH_4_Cl. After mixing these samples, relative quantitative information is achieved by comparing the intensity between the light and heavy isotopic peptide peaks. As the samples can be mixed at the beginning of sample processing, ^15^N metabolic labeling significantly reduces preparative and analytical variabilities, and allows extensive fractionation with little detrimental effect on quantification, enabling the relative quantification of thousands of proteins simultaneously (Arsova et al., 2012; Shrestha et al., 2022; Skirycz et al., 2011; Wang et al., 2002).

Our group has used this approach to identify and quantify immunoprecipitated plant Transport Protein Particle (TRAPP) complexes (Garcia et al., 2020) and ACINUS-mediated alternative splicing protein complexes (Bi et al., 2021), and affinity-purified BIN2 proximity-labeled protein interaction network (Kim et al., 2019). ^15^N labeling has also been applied to phosphoproteomics studies. For example, we identified and quantified S251 phosphorylation on BSU1 and demonstrated its flagellin-dependent manner (Park et al., 2019) and the Sussman lab has performed quantitative phosphoproteomics and identified RALF receptor FERONIA from Arabidopsis plasma membrane proteins (Haruta et al., 2014).

Despite its great promise, ^15^N labeling experiment has some drawbacks (Shrestha et al., 2022). As the data is typically acquired in data-dependent mode (DDA), missing values due to selecting different peptides for analysis in related runs is common. To ensure the most precursors are selected for fragmentation analysis in each run, DDA selects the most abundant ions observed in MS^1^ full scans for MS^2^ analysis. However, this will result in a different subset of peptides in the sample being measured in each experiment. Secondly, even with high-resolution data acquisition, co-eluting peptides or chemical noise in the MS^1^ scan still exist, especially in highly complex samples, resulting in inaccurate quantification. Thirdly, incomplete labeling in heavy-labeled samples leads to fewer proteins being identified, causing missing values among the different replicates (Minkoff et al., 2015; Shrestha et al., 2022). These drawbacks are more pronounced in lower abundance proteins where very few peptides are typically identified and quantified. Notably, these challenges are not all unique to ^15^N labeling quantification, yet better solutions are desired to improve the quantification quality.

Here we present utilizing Parallel Reaction Monitoring (PRM) (Peterson et al., 2012) in ^15^N labeling experiments to achieve more accurate quantification results (Bi et al., 2021; Park et al., 2019). In the PRM method, a pre-determined list of targeted peptide ions is mass filtered, usually using a quadrupole; they are fragmented, and all the yielded fragment ions are then detected, in our case, using a high-resolution Orbitrap mass analyzer (Figure 1). Because this is a targeted method, all peptides of interest are fragmented in reciprocal labeled replicates. Also, because the quantification is done using MS^2^ fragment ions measured in a high resolution, high accuracy instrument, signals from co-eluting peptides or chemical noise can be easily eliminated, leading to more accurate quantification.

**Figure 1:**
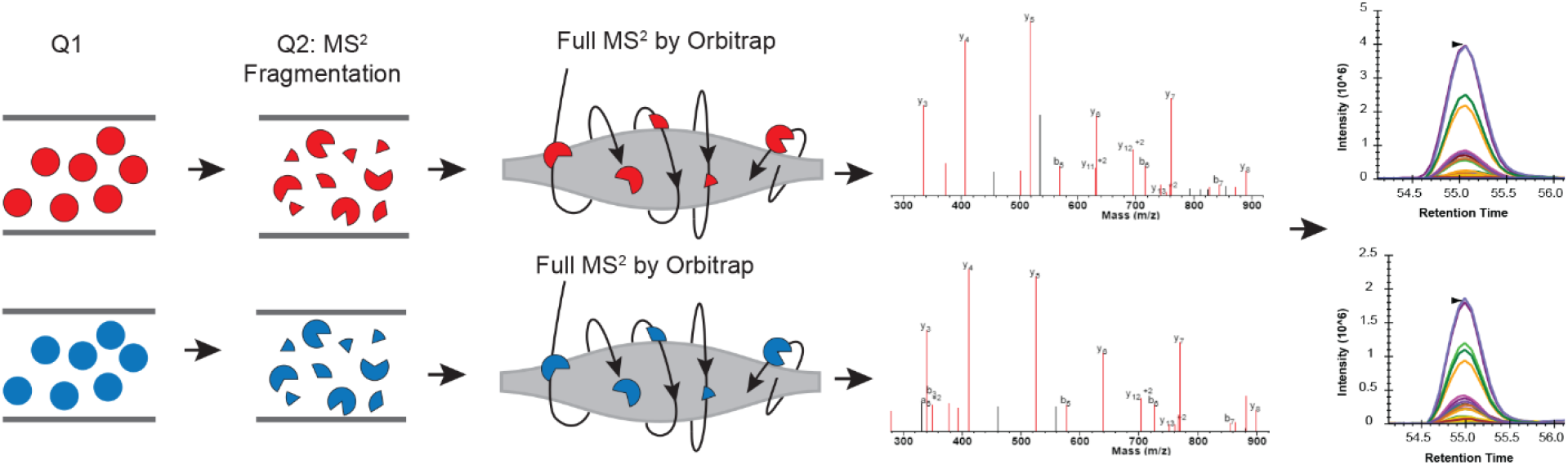
Schematic view of combining Parallel Reaction Monitoring to ^15^N metabolic labeling quantification workflow to get unambiguous quantification results. In each PRM cycle, light (^14^N) (colored in red) and heavy (^15^N) labeled peptide (colored in blue) ions from the target list are sequentially isolated in the first quadrupole (Q1). These ions are then fragmented using, for example, higher-energy collision dissociation (HCD), in the second quadrupole (Q2). The resulting MS^2^ fragments are separated and detected by the Orbitrap. The MS^2^ fragments from the light and heavy pairs are measured/scanned multiple times (cycles) across their elution profile. The area under the curve (AUC) of the fragment ions are integrated for quantification.

In this methods paper, we present the challenges of ^15^N quantification using MS^1^ scans of protein SAP18 that was acquired using data-dependent acquisition mode(Shrestha et al., 2022). We then show step-by-step how to remedy these challenges by setting up a PRM targeted method for the light ^14^N and heavy ^15^N target peptides using Skyline. We show how to set up a PRM acquisition method on an Orbitrap Q Exactive HF (the method would be almost identical on any quadruple-Orbitrap instrument such as Thermo Exploris or Tribrid instrument), and discuss the effects of different parameters such as the monitoring time window, numbers of peptides monitored at a particular retention time, cycle time and scan numbers across the chromatographic elution profile. Finally, we demonstrate using Skyline examples of how PRM provides unambiguous quantification for SAP18. We highlight the additional advantage of PRM on the light and heavy peptides that generate the same pattern of fragment ions which co-elute but with different mass. Instead of using synthetic peptides to validate the targeted results in most of label-free experiments (Picotti et al., 2009), the heavy labeled peptides in ^15^N labeling experiments can serve as natural synthetic peptides. This method is applicable to protein-level quantifications in ^15^N labeling samples and can also be applied to post-translational modified peptides with slight modifications (Bi et al., 2021; Park et al., 2019).

## 2.1 Materials and equipment

Data from two batches of samples are presented. In the first set, materials and data acquisition were detailed in (Bi et al., 2021), Briefly, the wild type (Col) and *acinus-2 pinin-1* plants were grown on Hoagland medium containing ^14^N or ^15^N (1.34 g/L Hogland’s No. 2 salt mixture without nitrogen, 6 g/L Phytoblend, and 1 g/L KNO_3_ or 1 g/L K^15^NO_3_ (Cambridge Isotope Laboratories), pH 5.8) for 14 days under the constant light condition at 21–22 °C on vertical plates. Proteins were extracted from six biological samples (one ^14^N-labeled Col sample −1, two of ^15^N-labeled Col samples −2, −3, one ^15^N-labeled *acinus-2 pinin-1* sample −4 and two of ^14^N-labeled *acinus-2 pinin-1 sample* −5 and −6) individually using SDS sample buffer and mixed as the following: one forward sample F1 (^14^N Col/^15^N *acinus-2 pinin-1*, Mix 1+4) and two reverse samples R1 and R2 (^14^N *acinus-2 pinin-1/^15^N* Col, Mix 2+5, Mix 3+6) and separated by the SDS-PAGE gel with a very short run (~3 cm). Two segments (upper part (U) ranging from the loading well to ~50 KD; lower part (L) ranging from ~50 KD to the dye front) were excised, trypsin digested, and analyzed by liquid chromatography-mass spectrometry (LC-MS) as described in (Bi et al., 2021; Shrestha et al., 2022)on a Q-Exactive HF instrument using 50 cm column ES803(50 cm x 75 μm ID, PepMap RSLC C18, 2 μm, ThermoFisher). For data-dependent acquisition, precursor scan was from mass-to-charge ratio (m/z) 375 to 1600 and the 20 most intense multiply charged precursors eluting at any given time were selected for fragmentation. Peptides were fragmented with higher-energy collision dissociation (HCD). The above untargeted acquisition data was analyzed using Protein Prospector Protein Prospector(Chalkley et al., 2005; Shrestha et al., 2022). The targeted method is described below in method section and analyzed by Skyline(Schilling et al., 2012).

The second batch of samples was generated using the same seedling growth conditions as described in (Bi et al., 2021). Detailed descriptions can be found in Supplemental Method. Briefly, ^14^N labeled Col and ^15^N labeled *acinus-2 pinin-1* were mixed about 1:1 ratio and data-dependent acquisition (DDA) and targeted quantification (PRM) data were acquired on Q-Exacive HF instrument with trapping column Acclaim PepMap 100 (75 uM x 2cm, C18, 3 μm, Thermo Fisher), then separated using analytical column Acclaim PepMap RSLC (75um x 25cm, C18, 2 μm, Thermo Fisher).

The following items are required for data analysis in this method.

1. Skyline software (free download from http://skyline.ms)
2. (Optional) Microsoft office, including Excel
3. (Optional) R studio

## 3. Method

### 3.1 Generate the target peptides list

To target proteins of interest (POI) for PRM analysis, it is necessary to get peptide information for those targets, including peptide sequence, *m/z* (mass to charge ratio), and retention time. This information can be retrieved either from prior ^15^N DDA data analysis or from publicly available databases. If from the latter, then a wider monitoring window is required since the retention time can shift due to different columns or gradients.

As peptides serve as a proxy to proteins, selecting target peptides that enable the best results is critical to yield the most accurate ratio that is truly representative of POI. Criteria for compatible target peptides:

- Peptide length: 6-25 amino acids long.
- Missed cleavages: peptides should have no missed cleavages. Using peptides containing missed enzyme cleavage sites is not ideal because if the digestion is more complete in one of the biological repeats, then the missed cleavage version may not be present.
- Not expected to be modified: Peptides containing methionines or tryptophans are generally not a good choice as they could become partly oxidized, splitting the signal between multiple masses.
- Intensity: Peptides from the same protein have different ionization efficiencies, so choosing a peptide that gave a strong signal when previously analyzed will give better quantification accuracy.
- Unique to the protein of interest.

Note: if there are multiple peptides to choose from, then following the criteria above will generate optimal results. If few peptides have been previously identified, then users are presented with a limited choice and may be forced to use the less ideal peptides for quantification.

### 3.2 Generate the Skyline template and export the information

After creating a list of target peptides, the next step is to create a Skyline template for them. This will serve as a method to calculate the masses of the light and heavy versions of the peptides and will be used downstream to import and quantify peptide ratios once the PRM data is acquired.

#### 3.2.1 Organize the peptide sequence and protein name into a two-column Excel file

**Table 1:**
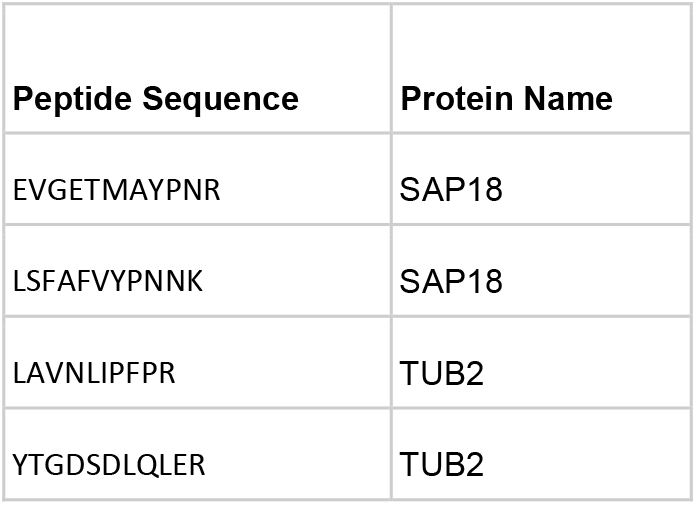
Example of Skyline peptide list.

#### 3.2.2 Import peptides into Skyline

Peptide templates in Skyline may be generated manually, using an R script or by using a library.

##### Manual template generation involves two steps

1) generate the light version; 2) modify the sequence to the heavy version with heavy isotope modification. Once modified, Skyline will automatically include the light version.

- To generate the light version, the most straightforward way is to copy the peptide sequences and protein names from an Excel spreadsheet like that shown in Table 1 and paste it into the peptides menu in Skyline. The peptides can be inserted into Skyline using the option:

Skyline > Edit > Insert > Peptides… > copy excel into skyline box ->insert.
- Modify the sequence to the heavy version with heavy isotope modification. Right-click on the peptide and click on edit modification. It is important to add ^15^N isotope heavy modification to Skyline. Detailed instructions are described in Supplemental Figure 1. Once all amino acids in the sequence have been modified with a heavy isotope, then the heavy version of the peptide is generated. Skyline will automatically include the light version.

##### Generate template using an R script

A faster way to insert target peptide pairs is by using an R script (available upon request) that can automatically generate heavy peptide sequences recognizable by Skyline. The sequence of the peptide is converted to a heavy version using the formula as listed in Table 2. The input file for the R script is the same as Table 1, and the output after running the script is a CSV file containing the heavy peptide sequences (Table 3). Insert the peptide sequence and protein name to Skyline as described above and Skyline will generate both light and heavy peptide templates.

**Table 2:** Light ^14^N to heavy ^15^N amino acid conversion for 20 amino acids.

| Amino Acid | Conversion |
| --- | --- |
| A,D,E,F,G,I,L,M,P,S,T,V,Y | [+1] |
| K,N,Q,W | [+2] |
| H | [+3] |
| R | [+4] |
| C + Carbamidomethylation | [+58] |
| M + oxidation | [+17] |

**Table 3:** Output from the R script that converts peptide sequence to heavy peptide sequence recognizable by Skyline.

| Peptide Sequence | Protein Name |
| --- | --- |
| E[+1]V[+1]G[+1]E[+1]T[+1]M[+1]A[+1]Y[+1]P[+1]N[+2]R[+4] | SAP18 |
| L[+1]S[+1]F[+1]A[+1]F[+1]V[+1]Y[+1]P[+1]N[+2]N[+2]K[+2] | SAP18 |
| L[+1]A[+1]V[+1]N[+2]L[+1]I[+1]P[+1]F[+1]P[+1]R[+4] | TUB2 |
| Y[+1]T[+1]G[+1]D[+1]S[+1]D[+1]L[+1]Q[+2]L[+1]E[+1]R[+4] | TUB2 |

**Table 4:** PRM template as an inclusion list.

| Mass [m/z] | Formula [M] | Formula type | Species | CS [z] | Polarity | Start [min] | End [min] | (N) CE | (N)CE type | MS X ID | Comment |
| --- | --- | --- | --- | --- | --- | --- | --- | --- | --- | --- | --- |
| 650.3402 |  |  |  | 2 | Positive | 87.35 | 107.35 | 27 | NCE |  | LSFAFVYPNNK |
| 657.3195 |  |  |  | 2 | Positive | 87.35 | 107.35 | 27 | NCE |  | L[+1]S[+1]F[+1]A[+1]F[+1]V[+1]Y[+1]P[+1]<br>N[+2]N[+2]K[+2] |
| 633.7928 |  |  |  | 2 | Positive | 36.43 | 56.43 | 27 | NCE |  | EVGETMAYPNR |
| 641.2706 |  |  |  | 2 | Positive | 36.43 | 56.43 | 27 | NCE |  | E[+1]V[+1]G[+1]E[+1]T[+1]M[+1]A[+1]Y[+1]<br>P[+1]N[+2]R[+4] |

##### Generate the template using a library

Templates can be generated using the library function. Identified peptide information is first exported from the search engine results and imported into Skyline. Detailed instructions can be found in Skyline tutorials (https://skyline.ms/_webdav/home/software/Skyline/%40files/tutorials/MethodEdit-2_5.pdf?listing=html) and will not be covered here. The advantage of using a library is that it allows the generation of templates for hundreds or more peptides and it can contain a retention time which potentially can be used for data filtering in the subsequent steps. The downside of using the library is that the light and heavy peptides are input as separate entries, creating potential problems for data analysis unless users manually curate the template using the steps outlined above.

#### 3.2.3 Export list containing mass information of paired peptides from Skyline

Once all the appropriate target peptides are represented in the Skyline session, the light and heavy lists are exported to help generate the spreadsheet needed for the PRM acquisition method in the Orbitrap mass spectrometer.

The path to export the skyline list is:

Skyline > File > Export > Report.

Export a custom report that only contains peptide sequence, precursor charge, protein name, and precursor m/z.

To create a custom report, navigate:

Edit List > Add.. > Check Peptide Sequence, Precursor Charge, Protein Name, and Precursor Mz

### 3.3 Add retention time and create PRM template (inclusion list)

#### 3.3.1 Add retention time window

A critical component of the PRM assay is the inclusion of a retention time window. From either a previous data-dependent acquisition MS run or from a public database, copy and paste the recorded retention time for those target peptides into the report file generated by Skyline with a header labeled retention time. Next add a monitoring window that specifically corresponds to the precursor survey retention time window. A narrow window (±5 min or less) can be chosen if the same chromatographic conditions in DDA are to be used for targeted quantification. Otherwise, a wider window is recommended to make certain that the target peptides are monitored. Notably, the numbers of the targets, cycle time, and the data points across the elution profile for the PRM method are detailed in (Gallien et al., 2014; Peterson et al., 2012; Rauniyar, 2015). Typically, peptides elute in a 15-60 second window, depending on the column used. However, retention times in liquid chromatograph can vary slightly between runs, therefore, the actual monitoring window is normally wider than the actual peptide elution profile. Fig.2 compares a 20 minute’ window (A) to a 5 minute’ window (C), illustrating the consequences on the increased number of monitored target peptides at a given time when a wider window is used (B, D).

**Figure 2:**
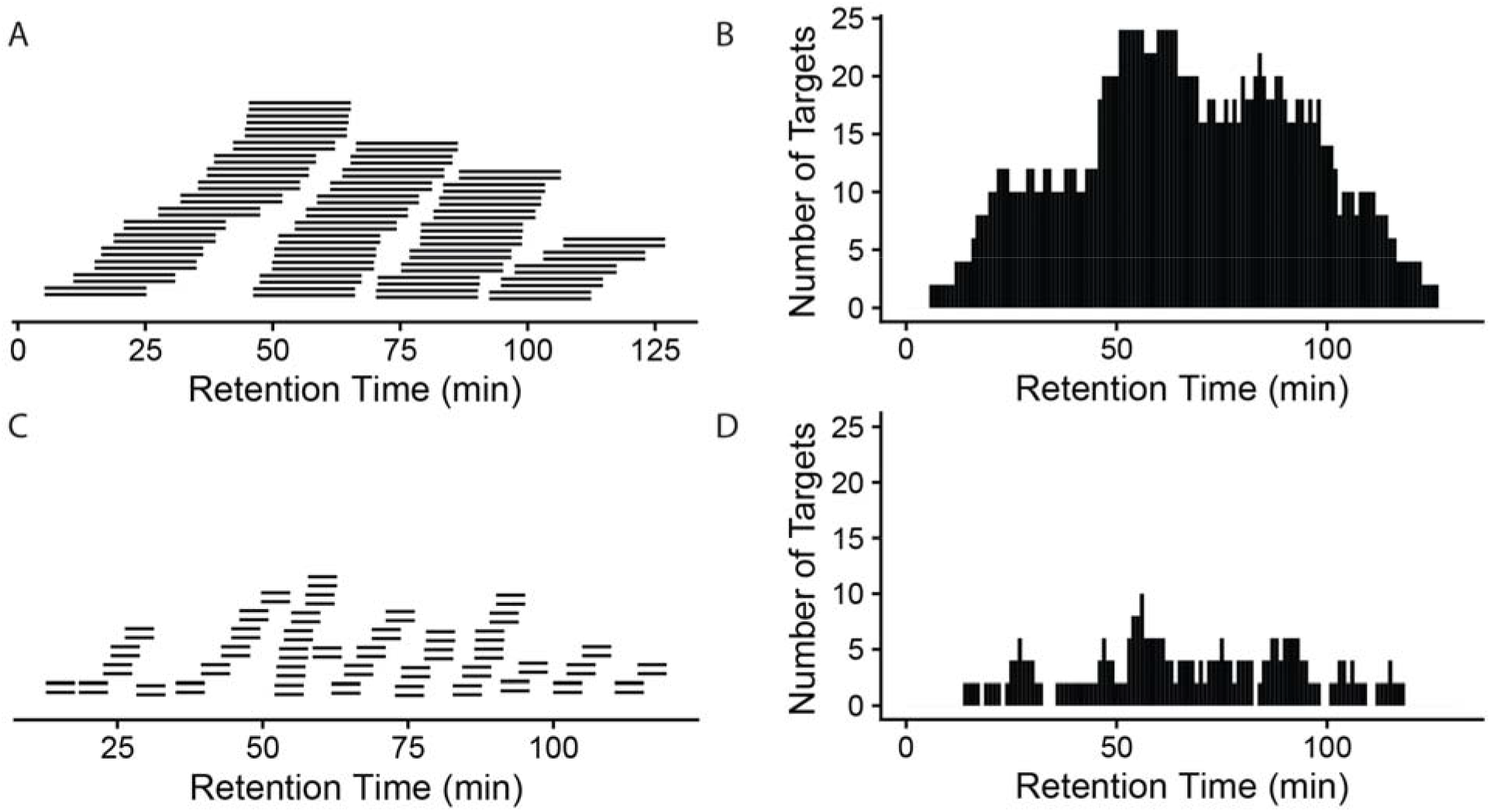
A narrow window is preferred to reduce monitored targets at a given time. A 20-minute window (A) will result in a longer target list (B) that will be monitored at a given time than a 5-minute window (C and D). x axis: retention time(gradient). A total of 42 peptide pairs (light and heavy) are in the targeted list across the whole gradient (A,C). More monitored targets in B will result in a longer cycle time (the time to cycle through the entire list of the target peptides at a given time on the gradient), meaning longer time between data points for a given peptide, so fewer measurements across the elution of the peptide.

The PRM cycle time will increase significantly when the target number increases in a cycle (Figure 3), and this will affect the number of data points across the chromatographic peaks. For reliable quantification, usually 8-10 scans across the peak are recommended (Gallien and Domon, 2015). If there are many precursors to target at a given time one may need to reduce the injection time (time accumulating a given precursor) to produce 8-10 data points across the peak. As a longer injection time is generally preferred to boost signal to noise, reduced injection time will impact sensitivity. Thus, target list, cycle time, and scan numbers (data points) need to be well balanced to achieve the best quantification results.

**Figure 3.**
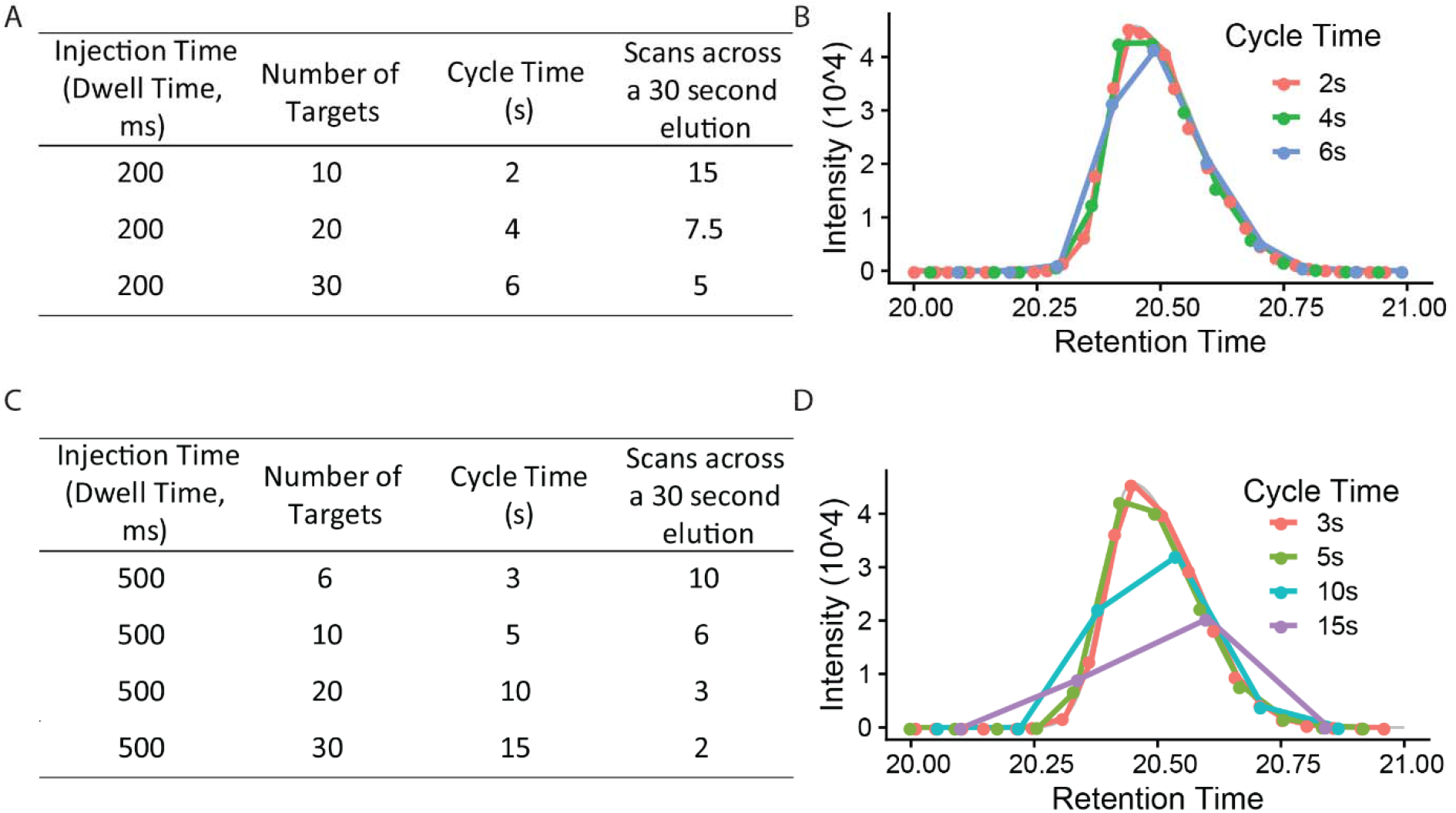
The relationship between injection time, number of targets, cycle time and scan number (points across a 30 second elution). (A) and (C) are using a 200 ms and 500 ms injection time each to calculate the cycle time and scan numbers (B, D). x axis is retention time (gradient, minutes). Injection time (the time to accumulate ions) is, in general, a rate limiting step in PRM method. 8-10 scans across the chromatography peak are preferred to give a good quantification result.

#### 3.3.2 Make a PRM template

This adapted Skyline report now has all the information needed to produce the form needed by the Orbitrap instrument to target these peptides. The next step is to create a csv file with the following headers, which can be imported into the PRM template via global list/inclusion list.

For a Q-Exactive HF, we recommend the “(N)CE” column should be “27” for all rows; the same as used in the DDA method. The first column numbers are generated from Skyline. Only the mass, polarity, retention time window, and collision energy information are required in the inclusion list. For other instruments, we recommend using the same collision energy used for DDA analysis in the PRM studies, as different collision energy can affect the fragment (transition) patterns (Diedrich et al., 2013).

### 3.4 Create PRM method on Orbitrap Analyzer

The PRM method is set up using PRM in the Q Exactive HF Orbitrap MS. Recommended parameters include MS^2^ resolution: 60,000, AGC target: 2e5, Maximum injection time IT: 200ms, Isolation window: 1.4 m/z, (N)CE: 27. Method runtime and chromatography conditions should be the same as used in DDA. Note: The Maxium injection time can be varied depending on the abundance of peptides and the number of peptides targeted at a particular retention time.

### 3.5 Import to Skyline, analyze the data, and export the result

Once the targeted data is acquired, it can be analyzed in the Skyline software.With the previously created peptide template in 3.2.2, we must first adjust the transition settings so that the raw data can be properly imported into Skyline (see Supplemental Figure 2). Raw data can be imported directly into Skyline using the option:

File > Import > Results >…

Once imported, quantification between the light and heavy version of each target peptide can be analyzed. Then we will export the results using the option:

File > Export >…

The following will be selected for the report, including: Precursor m/z, Best Retention Time, Peptide Sequence, Protein Name, Total Area Fragment, and File Name. The Total Area Fragment field contains the sum of the all the areas under the curve of the elution profiles of each transition ions.

## 4. Results

### Two common challenges in ^15^N quantification using DDA acquisition and quantification: missing values and contamination from co-eluting peptides

The goals of our discovery experiments were to understand: 1) Are *acinus-2* and *pinin-1* mutant null alleles? 2) Do SAP18 and SR45, two interactors of ACINUS and PININ, have less accumulation when ACINUS and PININ are absent? 3) What proteins are over accumulated or less accumulated in *acinus-2 pinin-1* double mutant? We performed three ^15^N metabolic labeling experiments, including one forward sample F1 (^14^N Col/^15^N *acinus-2 pinin-1*) and two reverse samples R1 and R2 (^14^N *acinus-2 pinin-1*/^15^N Col) experiments. Using a non-targeted DDA method we were able to quantify thousands of proteins and demonstrate ACINUS and SR45 proteins have dramatically reduced levels in the double mutant. We also identified proteins that are over-accumulated or less accumulated in the mutants (Shrestha et al., 2022). However, as is common in DDA data there were missing values in reciprocal labeled experiments, particularly for some of the lower abundance proteins. In the following, we used SAP18 as an example to demonstrate the challenge using DDA and how the use of targeted quantification improved the sensitivity and accuracy of quantification.

Using DDA, we were only able to identify and quantify SAP18 in two experiments (F1, R1), but not in the R2 experiment (Figure 4, A and B). In F1, we observed SAP18 had a dramatic reduction in the *acinus-2 pinin-1* double mutant. Four peptides were identified by MS/MS in F1 (all from the Col ^14^N channel), with three peptides showing consistent reductions in the mutant, such as the peptide shown in Figure 4C, while one peptide quantification showed an outlier result (Figure 4, A and D). The quantification ratio 0.386 of peptide “EVGETMAYPNR” was an outlier result because the matching peak of the ^15^N peptide was contaminated with a co-eluting ^14^N labeled peptide. We reason this is a co-eluting ^14^N labeled peptide based on the isotope pattern and that the M-1 peak is missing (labeled with black arrow), which we would expect to be about 73% of the intensity of the M peak based on the observed labeling efficiency of 94%. In contrast, in the reverse labeling experiment R1 in which Col was ^15^N labeled, SAP18 was only identified with a single peptide from Col, but quantification based on this peptide is wrong because the ^14^N peak isotope cluster is incorrect (Figure 4E) and the ^14^N monoisotopic peak is likely contaminated with chemical noise. In the reverse labeling experiment R2, no peptides were identified from SAP18 in both Col and *acinus-2 pinin-1*, hence no quantification was possible (Figure 4A and 4B).

**Figure 4:**
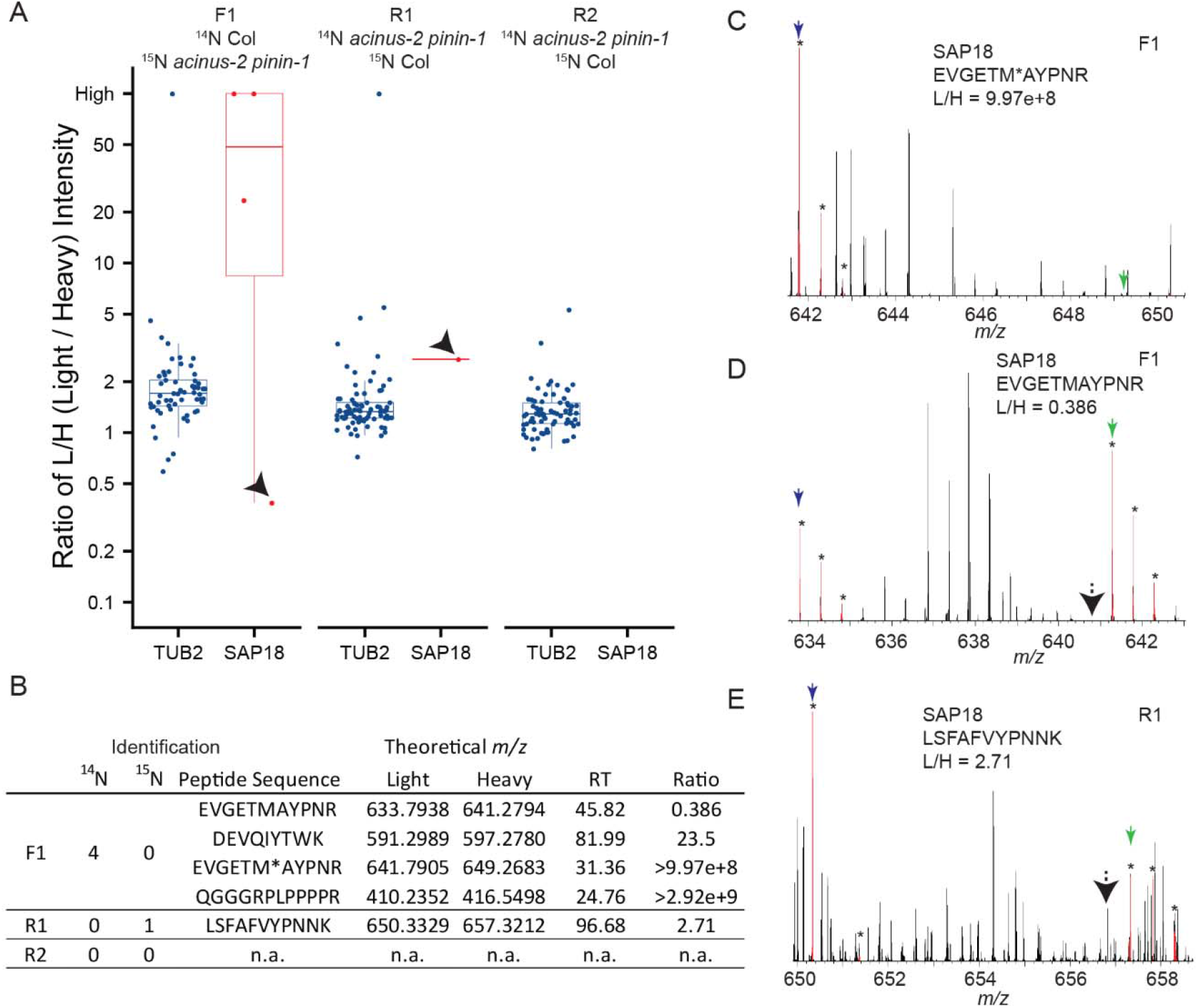
Quantification challenges in ^15^N metabolic labeling samples. (A and B) Boxplots of protein ratio measurements from reciprocal ^15^N labeling experiments: one forward sample F1 (^14^N Col/^15^N *acinus-2 pinin-1*) and two reverse samples R1 and R2 (^14^N *acinus-2 pinin-1*/^15^N Col). Each dot represents the ratio for a single peptide identified to each protein. TUBULIN2 is quantified in all three experiments based on many peptides, while SAP18 is quantified in two of three experiments (F1, R1), with inaccurate quantification in R1, and no quantification in R2. (C) Spectrum showing correct quantification of ^14^N and ^15^N pair for peptide “EVGETM(oxidation)AYPNR” from SAP18 protein in the F1 sample. (D) Quantification for peptide “EVGETMAYPNR” is inaccurate because the ^15^N matching peak is contaminated with a co-eluting peptide in F1 sample. (E) Quantification for peptide “LSFAFVYPNNK” is wrong as the matching ^14^N peak is contaminated with overlapping peaks. M, M+1, M+2 peaks labeled with star (*) are highlighted in red by Protein Prospector software; blue and green arrows point to ^14^N and ^15^N monoisotopic (M) peaks, respectively; black arrows in (D) and (E) point to M-1 peak position.

### Parallel Reaction Monitoring (PRM) on ^15^N metabolic samples

Targeted quantification using MS^2^ can address the challenges illustrated above. We have previously employed PRM to phosphopeptides in label-free samples (Ni et al., 2017) and in ^15^N samples (Park et al., 2019). Targeted quantification of ^15^N samples has also been implemented by the Sussman lab using selected reaction monitoring (SRM) using a low-resolution triple quadrupole mass spectrometer (Minkoff et al., 2015). Due to the high complexity of the samples because all proteins were extracted, PRM on a high resolution, high mass accuracy Orbitrap mass spectrometer should provide more accurate results. Two peptides from TUBLIN2 proteins (Figure 5), as well as 42 different peptide pairs (light and heavy) from 30 relatively abundant proteins in the samples (Supplemental Figure 3), were set up for PRM targeted quantification. From this data, we observed that: 1) ^14^N and ^15^N peptides almost co-elute, with ^15^N eluting about 2-4 seconds earlier in a 2-hour gradient (Supplemental Figure 3). 2) The light and heavy pair of peptides produce similar fragmentation patterns with different fragment masses, as illustrated in Figure 5 and Supplemental Figure 3. Each colored line represents the same fragment ion between the light and heavy pairs. 3) Peptides from the same protein give consistent quantification in PRM. The PRM measurement on TUB2 shows that samples were not mixed exactly at 1:1, similar to quantification result using MS^1^ level (Figure 4A). The ratios from TUB2 quantification were utilized for normalization for sample loading as the level of this protein is not expected to be altered in the mutants.

**Figure 5:**
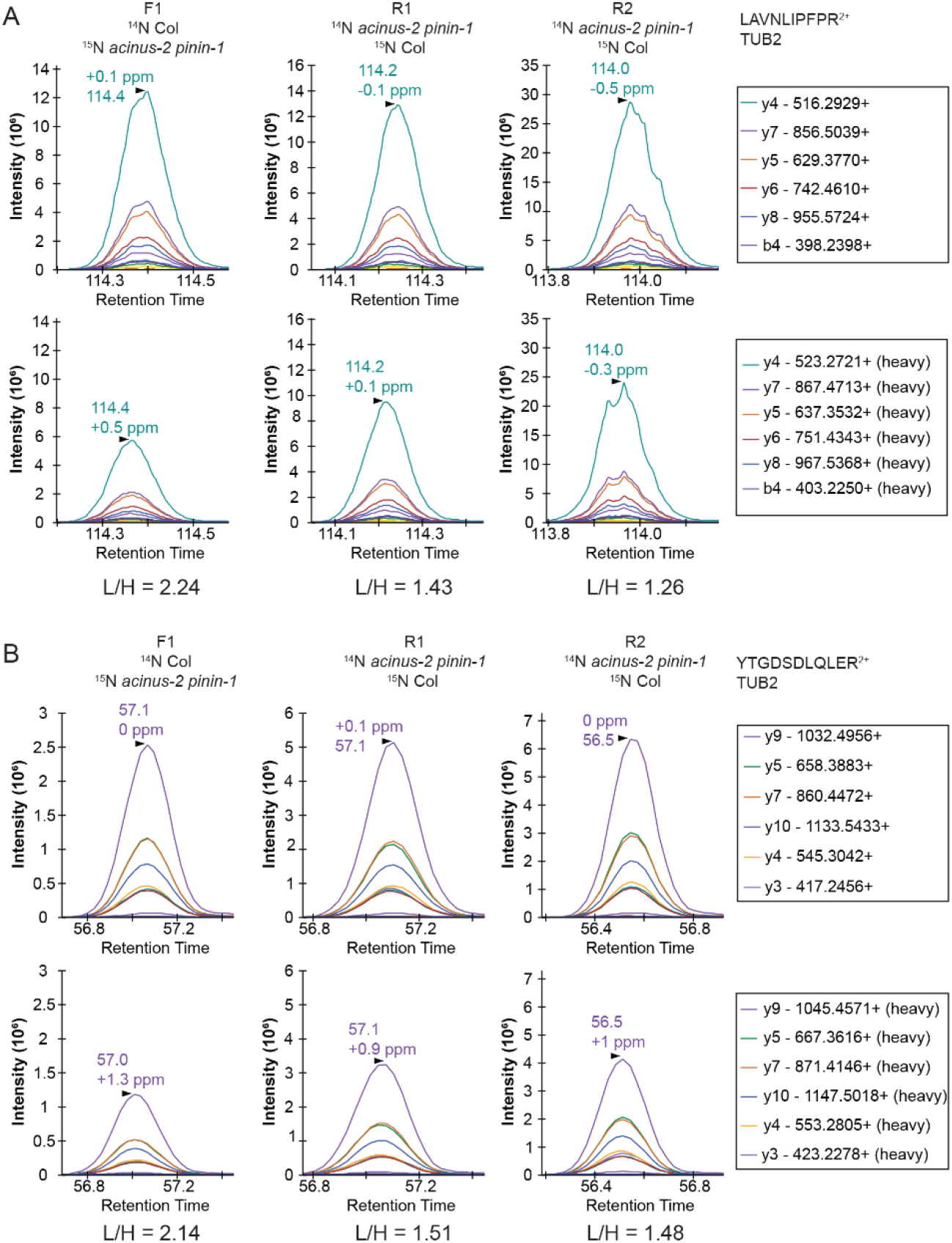
PRM quantification on TUBULIN2. Peptides “LAVNLIPFPR” (A) and “YTGDSDLQLER” (B) from TUB2 were quantified in three samples. The six most abundant fragment ions are annotated with mass in the right legend with the order from most to the least abundant. The light and heavy peptides generate the same pattern of fragment ions with different mass.

We set up targeted quantifications for the peptides “EVGETMAYPNR” and “LSFAFVYPNNK” from SAP18 as described in the Method section. The PRM quantification of “EVGETMAYPNR” showed a consistent reduction of this peptide in the *acinus-2 pinin* double mutant in all three replicates, confirming the ^15^N peak quantification in MS^1^ spectra was indeed incorrect due to coeluting peptide contamination (Figure 6A, Figure 4D). For the peptide “LSFAFVYPNNK”, the PRM quantification also showed a consistent reduction of signal in the double mutant (Figure 6B). More importantly, both peptides were quantified in the R2 sample, which was not quantified in the nontargeted analysis. We also performed targeted quantification of a third peptide “QGGGRPLPPPPR” (Bi et al., 2021). Altogether, we concluded SAP18 protein levels were dramatically reduced to ~2.7% of WT levels after normalization (median number of all the measured ratios), indicating that the stability of the other members of the ASAP and PSAP complexes is dependent on ACINUS and PININ in Arabidopsis.

**Figure 6:**
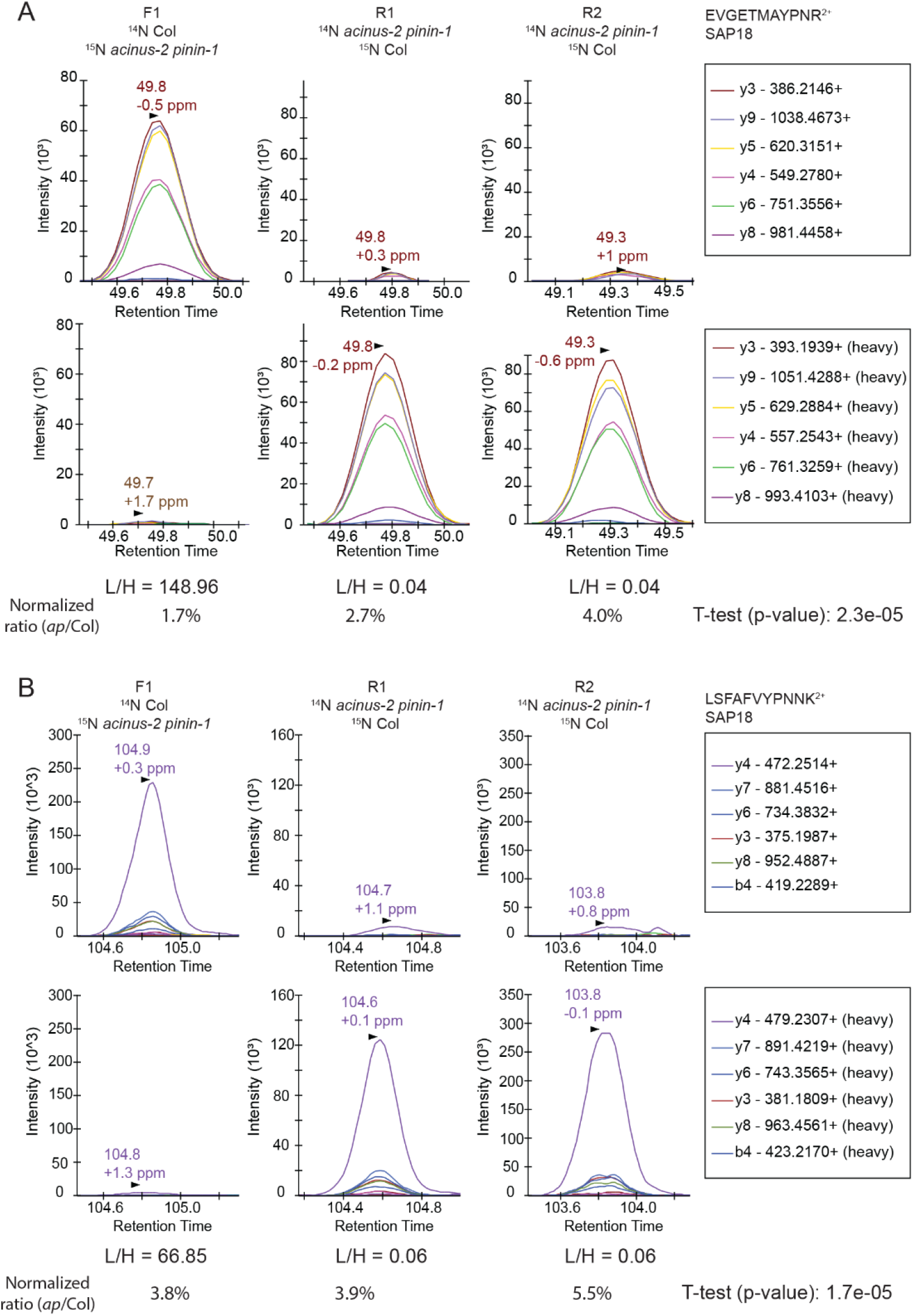
Targeted quantification using PRM on SAP18 proteins provides unambiguous quantification results. The targeted quantifications in three samples gave consistent results showing SAP18 was dramatically reduced in *acinus-2 pinin-1* mutant after normalization to TUB2. The light and heavy peptides gave a similar pattern of fragmentation. The peptides are quantified using area under the curve (AUC). (A) PRM results from peptide “EVGETMAYPNR” confirms the heavy matching peaks in Fig 4C is indeed from a co-eluting ^14^N labeled peptide; (B) PRM results from peptide “LSFAFVYPNNK” confirm the matching ^14^N peak in MS^1^ Fig 4E from DDA data is indeed contaminated with overlapping peaks.

Using the same approach, we also performed a targeted assay to prove *pinin-1* mutant was a null allele (Supplemental Figure 4). Notably, the PININ protein was of relatively low abundance and was not detected in DDA data. We were able to set up PRM for PININ using the retention time and peptide information obtained from previous immunoprecipitation results (unpublished). Using the PRM method, we showed both the N- and C-termini of PININ (before and after the T-DNA insertion) (Supplemental Figure 4) were detectable in Col but were not detectable in the double mutant (Bi et al., 2021).

## 5. Discussion

DDA is the most commonly used data acquisition approach in proteomics. It allows identification and quantification of many proteins without prior knowledge of the sample composition, so it is often used for the discovery stage (Shrestha et al., 2022). However, DDA can suffer from missing values for low abundance proteins due to its stochastic nature of selecting peaks for MS/MS. Furthermore, DDA of ^15^N heavy labeled samples often leads to less identifications and missing quantification values due errors in monoisotopic peak assignment caused by incomplete labeling (Shrestha et al., 2022). In addition, quantification using the MS^1^ scan often suffers from signal contamination from co-eluting peptides or chemical noise which will produce inaccurate ratios. To improve the quantification, we performed targeted quantification on these ^15^N labeled samples.

Targeted acquisition of ^15^N samples has previously been reported (Hart-Smith et al., 2017), where an inclusion list for proteins of interest was added to data acquisition, such that more proteins were repeatedly identified and quantified across multiple runs. However, the quantification was done at the MS^1^ level, so although this addresses the missing value problem, the quantification accuracy can still be affected by contaminating co-eluting peaks.

We demonstrated using PRM targeted quantification we were able to consistently quantify SAP18 in all replicates. SAP18 is not only of low abundance but it is also relatively small (18 KDa) so it produces few peptides, both of which contribute to the challenge of quantifying this protein. We were also able to quantify PININ proteins using PRM targeted assay even PININ proteins were not detected by DDA.

For PRM analysis at initial testing, it is recommended that users start with 5 peptides, as suggested by (Picotti et al., 2009). After testing, the two best peptides (combination of high ionization efficiency and multiple transitions) are chosen for final analysis. Employing PRM measurements on high resolution high accuracy instruments produces quantification results with low contamination from background. Sometimes there is interference in one or two transitions, but this can be easily spotted due to it being unlikely that the contaminating signal perfectly coelutes with the other transitions from the peptide of interest, so one can simply not use the problematic transitions for the quantification of that peptide. This is one of the advantages of PRM over SRM, as the full MS^2^ of the targeted peptides is measured, thus users can afford to remove one or two contaminated transitions without compromising the final quantifications results.

For reliable quantification, it is recommended to quantify based on at least 3 intense transitions and using more is typically better. However, some peptides (particularly those containing prolines) can produce one or two dominant transitions so sometimes a smaller number must be used. Fragment ions with an *m/z* above the precursor m/z are desirable as they are more specific. However, these are often not the most intense transitions, and so selecting only these transitions can decrease the sensitivity.

Combining PRM and ^15^N labeling has additional advantages. As shown in Figure 6, SAP18 is quantified using both ^14^N labeled and ^15^N labeled peptides that co-elute, and as MS^2^ of these pairs produce the same fragments with the same pattern (the order of most abundant to the least) but with different masses, the false-discovery rate of these peptides are minimal, thus the quantification of the proteins is even more reliable than using label-free samples. Many practitioners of the PRM experiments include synthetic (heavy) peptides to validate targeted quantification results (Gallien and Domon, 2015; Picotti et al., 2009). The ^15^N labeled heavy peptides in our experiments can serve as the same role as “natural” heavy peptides to provide this extra level of reliability.

## Data Availability Statement

The dataset for this study has been deposited in PRIDE. PRM data on TUB2 and SAP18 are available via ProteomeXchange with identifier PXD030096. DDA data on 42 peptide pairs are available with identifier PXD030098. PRM data on the 42 peptide pairs are available with identifier PXD030112.

## Author Contributions

A.V.R., R.S., and S.-L.X. designed the experiments, and A.V.R and R.S. performed the experiment, A.V.R., R.S., and S.-L.X. analyzed the data sets and generated figures. P.R.B. and R.J.C. provided technical support for Protein Prospector and revised the manuscript. A.V.R., R.S., and S.-L.X. wrote the manuscript.

## Funding

This research was funded by the NIH grant R01GM135706 to S.-L.X. and Carnegie endowment fund to Carnegie mass spectrometry facility.

## Conflict of interest

The authors declare that the research was conducted in the absence of any commercial or financial relationships that could be construed as a potential conflict of interest.

## Supplemental Material

The Supplemental Figures 1–6 and the supplemental method are provided.

**Supplemental Figure 1.**
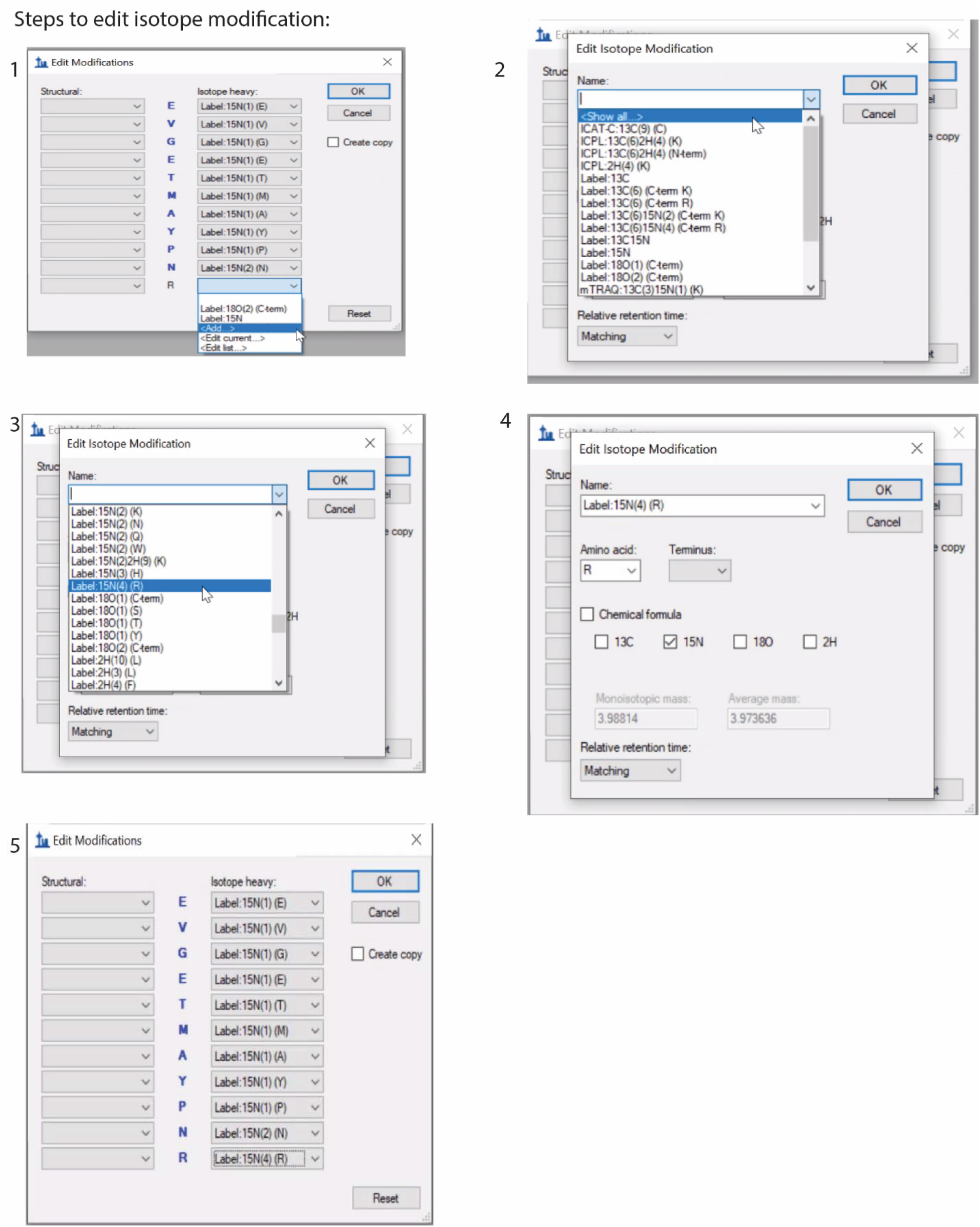
Example of steps for adding heavy isotope modification.

**Supplemental Figure 2.**
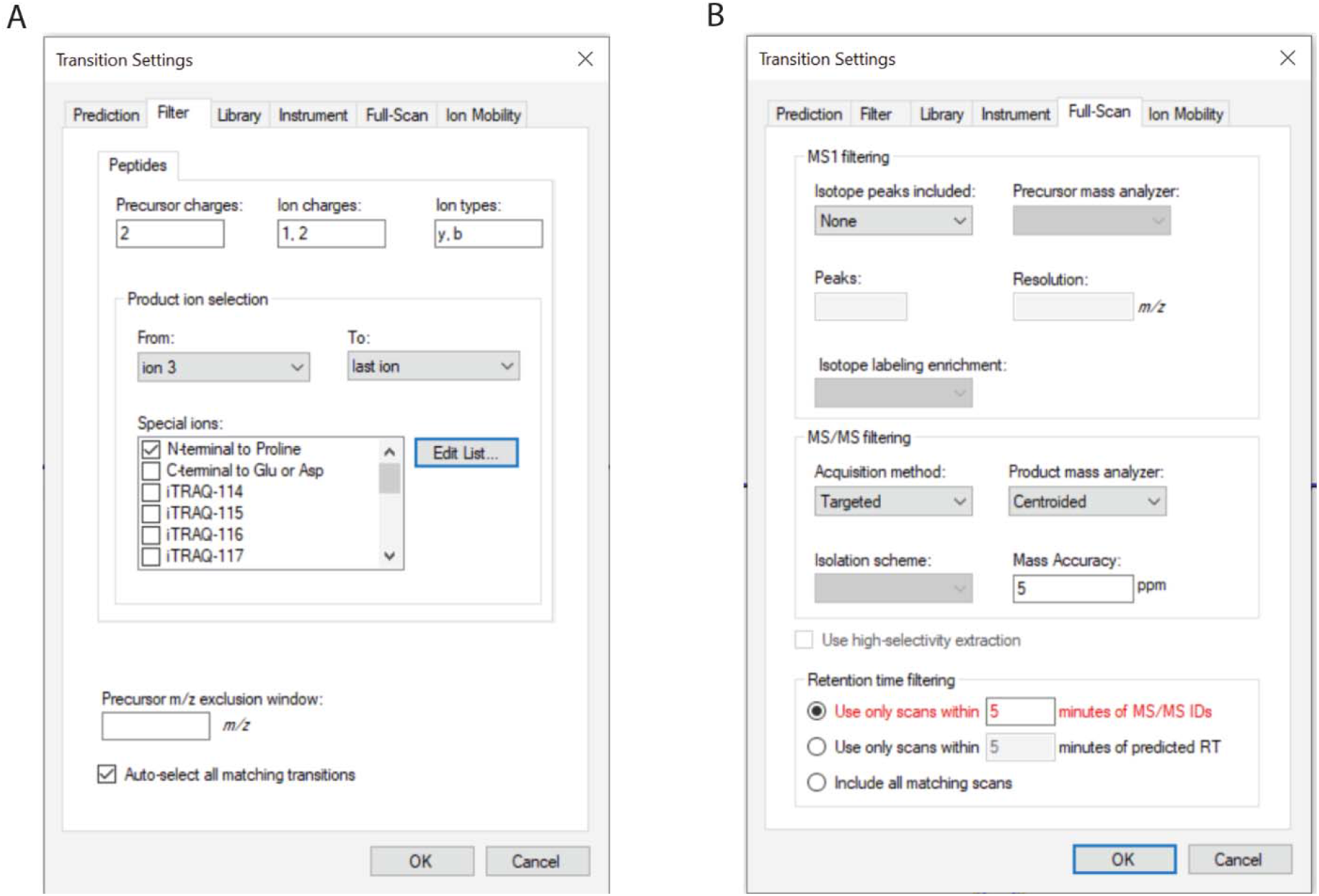
Example of setting up transition setting.

**Supplemental Figure 3.**
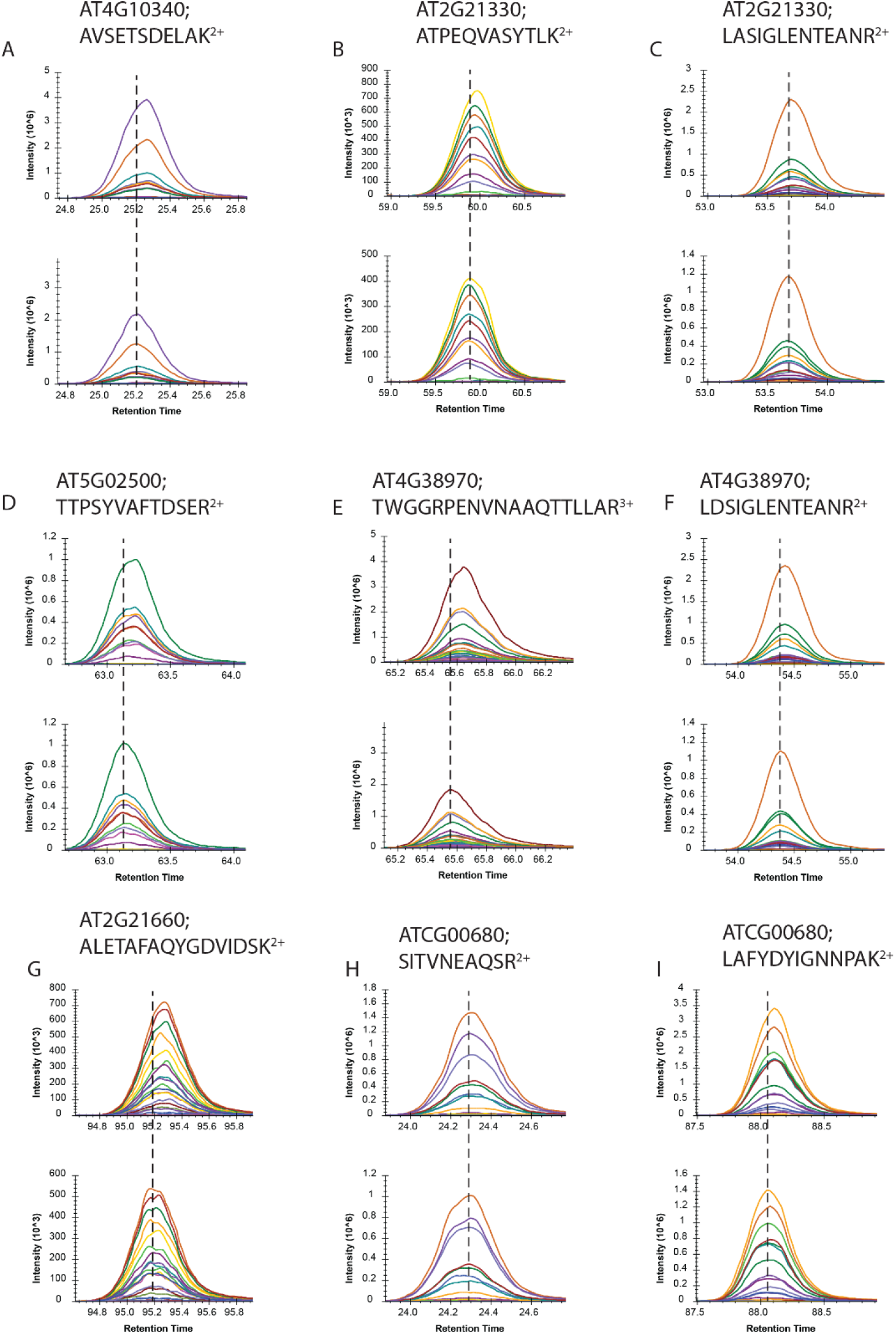
PRM quantification on multiple proteins. 42 pairs of peptides (^14^N and ^15^N peptides) are quantified using the ^14^N Col/^15^N *acinus-2 pinin-1* sample and 9 pairs are displayed. The top chromatograph represents ^14^N light fragment ions, and the bottom represents the ^15^N heavy fragment ions. The apex of the heavy peak is centered with the dashed line. The light and heavy peptide almost co-elute, with the heavy version elutes about 2-4 seconds earlier than the light version.

**Supplemental Figure 4.**
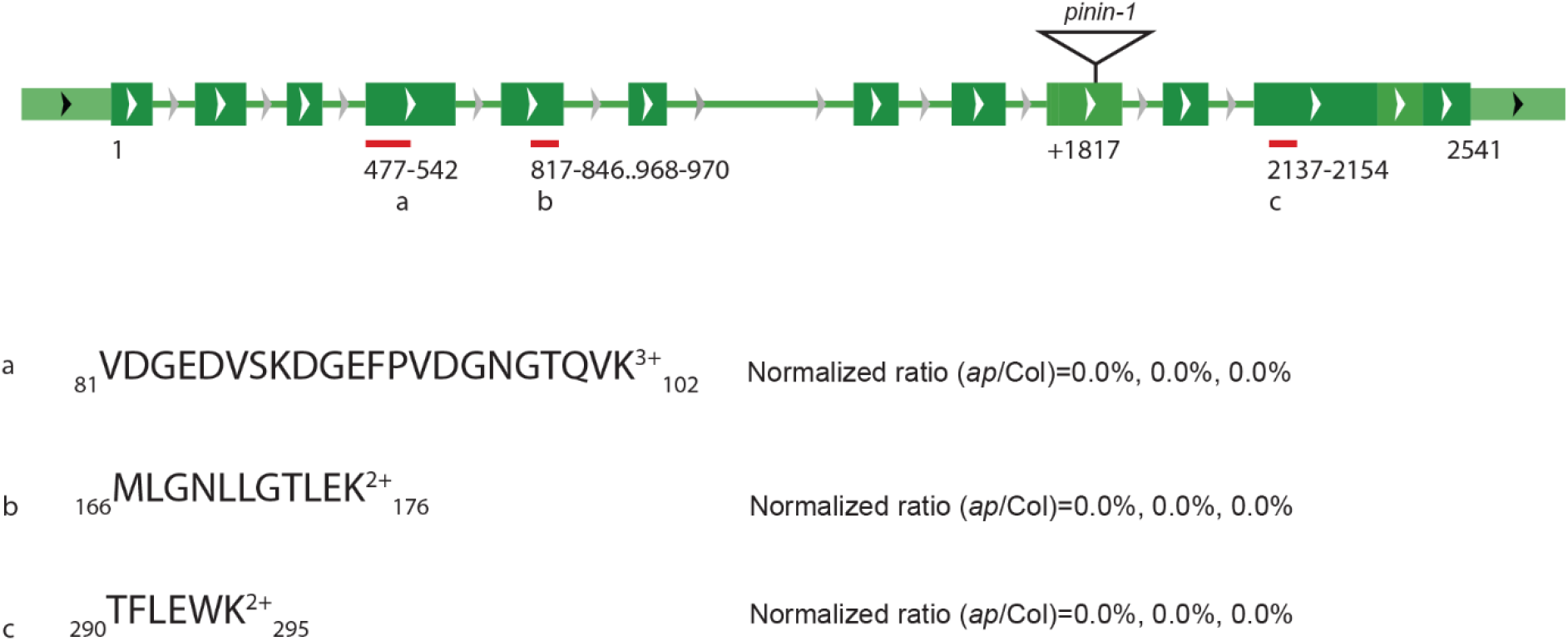
Targeted quantification using PRM on PININ proteins show *pinin* mutant is a null allele. Two peptides and one peptide in regions before and after the T-DNA insertion were used for the targeted analysis. PRM data shows both the N and C-termini of PININ were detectable in Col but not detectable in the *acinus-2 pinin* double mutant.

## Supplemental Method Method used for the second-batch sample preparation and data acquistion

The WT and *acinus-2 pinin-1* plants were grown on Hoagland medium containing ^14^N or ^15^N (1.34 g/L Hogland’s No. 2 salt mixture without nitrogen, 6 g/L Phytoblend, and 1 g/L KNO_3_ or 1 g/L K^15^NO_3_ (Cambridge Isotope Laboratories), pH 5.8) as described as (Bi et al., 2021). Proteins were extracted from two samples (one ^14^N-labeled Col, one ^15^N-labeled *acinus-2 pinin-1*) followed with protocols as described (Xu et al., 2017) with slight modification. Briefly, samples were first extracted individually using SDS sample buffer (0.1 M Tris·HCl, pH 8.0; 2% (wt/vol) SDS; 20 mM EGTA; 20 mM EDTA; 1.2% (vol/vol) Triton X-100; 2x protease inhibitor), then the protein concentration of each sample was measured by BCA assay kit (Thermo Fisher), subsequently followed by mixture to have 1:1 protein concentration mixture. Then Proteins were further extracted by cold phenol extraction. Proteins were digested with trypsin and the resulting peptides were de-salted using Sep-Pak waters C18 centrifuge columns. The peptides were analyzed on a Q-Exactive HF mass spectrometer (Thermo Fisher) equipped with an Easy LC 1200 UPLC liquid chromatography system (Thermo Fisher).

### Data dependent Acquisition

Peptides were first trapped using trapping column Acclaim PepMap 100 (75 uM x 2cm, nanoViper 2Pk, C18, 3 μm, 100A), then separated using analytical column Acclaim PepMap RSLC (75um x25cm, nanoViper, C18, 2 μm, 100A) (Thermo Fisher). The flow rate was 300 nL/min, and a 120-min gradient was used. Peptides were eluted by a gradient from 3 to 28% solvent B (80% (v/v) acetonitrile/0.1% (v/v) formic acid) over 100 min and from 28 to 44% solvent B over 20 min, followed by a short wash at 90% solvent B. For DDA acquisition, the precursor scan was from mass-to-charge ratio (m/z) 375 to 1600 and top 20 most intense multiply charged precursors were selected for fragmentation. Peptides were fragmented with higher-energy collision dissociation (HCD) with normalized collision energy (NCE) 27. DDA data was analyzed as described in (Shrestha et al., 2021).

#### PRM Targeted quantification

The data-dependent acquisition was used first to get the peptide information from multiple proteins with peptide mass/charge (m/z), retention time, and MS2 fragments. 42 paired peptides were picked that span the whole elution profile of 130 minutes. For targeted analysis, parallel reaction monitoring (PRM) acquisition using a 10-min window (±5 min) was scheduled with an orbitrap resolution at 60,000, AGC value 2e5, and a maximum fill time of 60 ms. The isolation window for each precursor was set at 2.0 m/z unit. PRM data was processed with a 5-p.p.m. window using skyline from ^14^N- and ^15^N-labeled samples.

## Notes

### Competing Interest Statement

The authors have declared no competing interest.

### Summary of Updates

We have added two new figures and revised the manuscript.

